# A set of endogenous control genes for use in quantitative real-time PCR experiments reveal that the wild-type formin *Ldia2* is enriched in the early pond snail embryo

**DOI:** 10.1101/660381

**Authors:** Harriet F. Johnson, Angus Davison

## Abstract

Although the pond snail *Lymnaea stagnalis* is an emerging model organism for molecular studies in a wide variety of fields including development, biomineralisation and neurophysiology, there are a limited number of verified endogenous control genes for use in quantitative real-time PCR (qRT-PCR). As part of larger study on snail chirality or left-right asymmetry, we wished to assay relative gene expression in pond snail embryos, so we evaluated six new candidate control genes, by comparing their expression in three tissues (ovotestis, foot, and embryo) and across three programs (geNorm, Normfinder and Bestkeeper). The specific utility of these control genes was then tested by investigating the relative expression of six experimental transcripts, including the formin *Ldia2*, a gene that has been associated with chirality in *L. stagnalis*. All six control genes were found to be suitable for use. Of the six experimental genes that were tested, it was found that all were relatively depleted in the early embryo compared with other tissues, except the formin gene *Ldia2*. Instead, transcripts of the wild type *Ldia2*^*dex*^ were enriched in the embryo, whereas a non-functional frameshifted version *Ldia2*^*sin*^ was severely depleted. These differences in *Ldia2*^*sin*^ expression were less evident in the ovotestis and not evident in the foot tissue, suggesting that nonsense-mediated decay may be obscured in actively transcribing tissues. This work therefore provides a set of control genes that may be useful to the wider community, and shows how they may be used to assay differences in expression in the early embryo.

## INTRODUCTION

The pond snail *Lymnaea stagnalis* is a hermaphrodite, pulmonate snail which is increasingly used in a wide range of research areas including ecology, evolution, development, neuroscience, behaviour, parasitology and sexual selection. Due to the species perhaps predominant prior use as a model system in neuroscientific studies, many of the earlier published molecular studies were confined to the central nervous system (e.g. Feng *et al.*, 2009). More recently, molecular studies have come from different fields, including especially ecotoxicology and biomineralisation (Bouetard *et al.*, 2012; Hohagen & Jackson, 2013). An unannotated draft genome sequence is available (Davison *et al.*, 2016), and there is a collaborative effort underway to produce a publically available, high-quality, annotated genome sequence (Genoscope-CEA, de la Recherche à l’Industrie, France).

In the past few years, *L. stagnalis* snails have also become an important organism in the study of left-right asymmetry, because the species exhibits genetically tractable variation in chirality (Kuroda *et al.*, 2009; Shibazaki, Shimizu & Kuroda, 2004). This recently culminated in the finding that a disabling mutation in one copy of a diaphanous-related formin *Ldia2* is associated with early symmetry breaking in the developing embryo (Davison*, et al.*, 2016). In preparing that work, we decided that it was necessary to design a new set of control genes to use with quantitative real-time PCR (qRT-PCR) in *L. stagnalis*.

Specifically, as we set out to measure the expression of cytoskeletal genes in *L. stagnalis*, this precluded the use of actin and tubulin as appropriate endogenous controls, because they are themselves cytoskeletal genes. Unfortunately, many of the previously published qRT-PCR studies on *L. stagnalis* either used ribosomal RNA (rRNA), actin or tubulin genes as endogenous controls (Bavan *et al.*, 2012; Bouetard *et al.*, 2013; Carter *et al.*, 2015; Hatakeyama *et al.*, 2013; Lu & Feng, 2011; Ribeiro *et al.*, 2010; van Kesteren *et al.*, 2006; van Nierop *et al.*, 2006). rRNA genes are potentially problematic because the over-abundance of rRNAs relative to the target mRNA sequence can lead to problems in accurate normalisation, and in any case, rRNA is transcribed through an independent pathway from mRNA and therefore not regulated in the same manner (Radonic *et al.*, 2004). More generally, it is now widely accepted that there are no universal endogenous control genes, and each gene intended for use as an endogenous control should ideally be validated as consistently expressed across all experimental conditions.

Therefore, we aimed to develop and test a new set of endogenous genes as controls, for use in our study, but also for subsequent use by the wider community, just as has been the case in some other species (Hibbeler, Scharsack & Becker, 2008; Li *et al.*, 2017; Olias *et al.*, 2014; Sirakov *et al.*, 2009).

## METHODS

### Sample preparation

Three separate tissues, single-cell embryo, ovotestis (hermaphrodite gonad) and foot, were used, all from laboratory reared individuals of *L. stagnalis*. Total RNA was extracted from embryos using the RNeasy micro kit (Qiagen), including DNAse treatment, yielding approximately 0.5 ng total RNA per embryo. Ovotestis and foot tissue samples were removed from individual adult snails and snap frozen using a dry ice/ethanol slurry. Total RNA was immediately extracted from them using TRI Reagent® solution (Applied Biosystems). Complementary DNA (cDNA) was then synthesised from a maximum of 500 ng total RNA, using Superscript III reverse transcriptase (Invitrogen) and random primer mix (NEB). Aliquots were then made of the experimental working concentration dilutions of cDNA to reduce freeze-thaw cycles, whereas serial dilutions were performed independently for each standard curve experiment. All cDNA samples were gently vortexed before use and prior to each serial dilution step.

### Primer design

Using transcriptomic resources of 1-2 cell stage *L. stagnalis* embryos (Liu *et al.*, 2014), six genes were selected as potential endogenous controls, all with well-characterised gene function. These included short-chain specific acyl-CoA dehydrogenase (*Lacads*); elongation factor 1-alpha (*Lef1a*); histone protein, H2A (*Lhis2a*); 60S ribosomal protein L14 (*Lrpl14*); ubiquitin-conjugating enzyme E2 (*Lube2*); and 14-3-3 protein zeta (*Lywhaz*). Primer pairs were then designed using Primer 3 (Rozen & Skaletsky, 2006), aiming for a Tm range within 2°C, and amplicon product sizes between 110-130bp, including GC clamps where possible, and making them exon-spanning and/or exon-crossing, to minimise problems with accidental genomic DNA carry over. To initially verify the primers, a standard PCR was used alongside a genomic DNA control sample and the products visualised on an agarose gel. Additionally, the specificity of the amplicons of all six primer pairs was verified through Sanger sequencing.

### Primer specificity and amplification efficiency

Primer efficiencies for each primer pair were calculated via standard curve qRT-PCR experiments using the Applied Biosystems 7500 fast system v2.3 and the same cycling parameters (below). Five standardised concentrations were used with an additional negative control (PCR grade water). Five-step serial cDNA dilutions were performed using molecular grade water and a dilution factor of 1:5. Primer efficiencies for all six endogenous control gene primer pairs were estimated using the same reference sample, created from pooling cDNA samples. Average primer efficiencies for each primer pair were then calculated via the arithmetic mean of a minimum of two successful standard curve experiments. A standard curve experiment was considered successful if it produced a R^2^ value of >0.98. Values from the lowest concentration dilutions were omitted if they dramatically reduced the amplification efficiency or R^2^ value of an experiment. The range of dilutions included in the standard curve experiment indicates the limits of acceptable working concentration/dilution factor for an experimental comparative qRT-PCR assessment.

Cycle threshold (Cq) values were obtained from qRT-PCR experiments using the ABI 7500 fast system v2.3. Each reaction contained 5 µl of Primer Design’s fast SYBR® green master mix, 0.5 µl forward and reverse primer (4 µM), 1.5 µl PCR grade water and 3 µl of cDNA. All samples were used at a 1:30 dilution of the original cDNA concentration. Mastermixes were prepared for each target gene experiment and a temperature melt curve step was included at the end of all qRT-PCR reactions. Thermocycling parameters were as follows: 95°C for 20, 95°C for 3 seconds, 60°C for 30s (data collection, Cq), 95°C for 15 seconds, cycle 39 more times, 60°C for 60 seconds, slow temperature ramp 1% (data collection; temperature melt curve), 95°C for 15 seconds, 60°C 15 seconds. All experimental samples were performed in triplicate repeat and negative controls in duplicate repeat for each of the six reference genes.

### Normalising control software

Three methods were used to assess the same qRT-PCR data, all of which run as macros within Microsoft Excel 2003. BestKeeper used raw Cq values (Pfaffl *et al.*, 2004), whereas NormFinder (Andersen, Jensen & Orntoft, 2004) and geNorm (Vandesompele *et al.*, 2002) required linearised Cq values. Efficiency-corrected linearised relative Cq values were calculated for each sample using the Pfafll method (Hellemans *et al.*, 2007). BestKeeper ran entirely from raw Cq values and corrected for amplification efficiency via the inbuilt formulas within the macro, via the manually-input amplification efficiency values.

### Relative expression of cytoskeletal genes

Variation in the left-right asymmetry of snails, or chirality, is under the control of a single maternally expressed locus. In *L. stagnalis*, maternal *D* alleles dominantly determine a clockwise (“dextral”) twist in offspring. For our experiments we created a single near-isogenic line of snails (>99%) that was still variable for the chirality locus, by repeated backcrossing (Davison*, et al.*, 2016). From this line, separate homozygous dextral (*DD*) and sinistral (*dd*) lines were produced, subsequently deriving heterozygote (*Dd*) snails by crossing individuals from the near-isogenic lines.

Previously, we reported finding that tandemly duplicated, diaphanous-related formin genes, *Ldia1* and *Ldia2*, are perfectly associated with variation in chirality of the pond snail, and that the sinistral-derived version of *Ldia2* contains a disabling frameshift mutation, which results in much reduced levels of *Ldia2* mRNA in the embryo (Davison*, et al.*, 2016). To further explore changes in expression between genotypes (*DD*, *Dd*, *dd*) and tissues (single-cell embryo, foot, ovotestis), the relative expression of *Ldia1* and *Ldia2* and four other genes was tested against the validated endogenous control genes. As above, primer pairs were designed using Primer 3 with the same conditions (Table 1). The other genes included the cytoskeletal gene furry *Lfry*, another tightly linked cytoskeletal gene, a fat-like cadherin *Lfat*, both tightly linked to *Ldia1/2* on the same chromosome, as well as unlinked actin-related proteins subunits 1a and 3, *Larp2/3-1a* and *Larp2/3-3*.

**Table 1.**
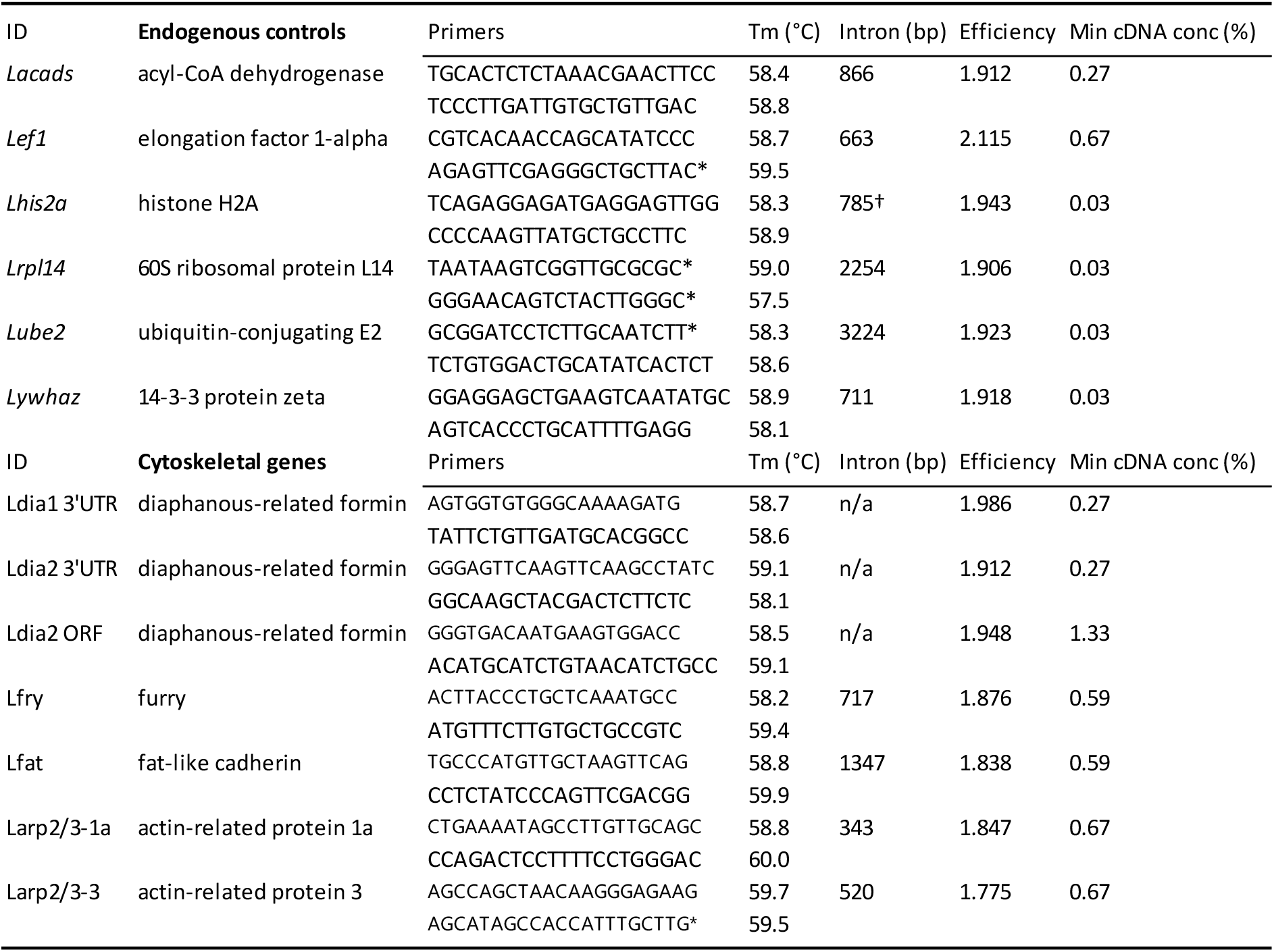
Primers used for the amplification of endogenous control genes (top) and the tested cytoskeletal genes (bottom), including the estimated intron size and the minimum concentration of sample cDNA (as a percentage of full concentration) required to achieve the amplification efficiency. *primer on exon/intron boundary.

Note that because of the high sequence identity between *Ldia1* and *Ldia2*, it was not possible to design intron-spanning PCRs for these loci. Instead, primer pairs were designed in the 3’UTR, because this region was most variable between copies, *Ldia1 3’UTR* and *Ldia2 3’UTR*, in addition to a primer pair in the open reading frame, *Ldia2 ORF*.

Relative expression of these genes was tested against the endogenous controls *Lhis2a*, *Lube2* and *Lywhaz* (embryo/foot) or *Lhis2a*, *Lube2* and *Lrpl14* (ovotestis), by calculating the Normalised Relative Quantity (NRQ) values from the average Cq value of each sample using the Pfaffl method (Hellemans*, et al.*, 2007; Pfaffl, 2001; Pfaffl*, et al.*, 2004). For each sample, first the relative quantity per target gene (ΔCq target) was calculated by subtracting the average Cq value of the sample from that of the calibrator sample. This ΔCq value was then corrected for amplification efficiency (E) by multiplying ΔCq to the base percentage amplification efficiency (represented as a value between 1 and 2). The efficiency-corrected relative quantities were then normalised to the endogenous control genes by dividing by the geometric mean (geoM) of the efficiency corrected delta Cq values calculated for each of the control genes (ΔCq ref) in the same manner as described above.

## RESULTS

### Primer specificity and amplification efficiency

All control primer pairs demonstrated amplification efficiencies between 1.906 and 2.115 with R^2^ values exceeding 0.98. All primers demonstrated acceptable amplification efficiencies in dilutions of 1:150 (0.67%), or less, of the full concentration. The working concentration of a 1:30 dilution used in the subsequent qRT-PCR experiments fell well within these limits.

### Comparing normalising control software

In the embryo, geNorm placed *Lhis2a* and *Lube2* as the most stable pair of genes, with a combined stability score of 0.196 (Table 2). The inclusion of any number of the genes provided a V score of <0.15, indicating that the combination of genes will provide a reliable normalisation factor (PrimerDesign 2014). The lowest V score was achieved with the inclusion of the five genes *Lhis2a*, *Lube2*, *Lrpl14*, *Lacads* and *Lywhaz*. In the foot, geNorm placed *Lywhaz* and *Lube2* as the most stable pair of genes with a combined stability score of 0.217. The inclusion of any number of the genes provided a V score of <0.15, although the lowest V score was achieved with the inclusion of the four genes *Lywhaz*, *Lube2*, *Lhis2a* and *Lacads*. In the embryo, geNorm placed *Lrpl14* and *Lube2* as the most stable pair of control genes with a combined score of 0.250. *Lhis2a* bore the lowest M score of all the target genes, at 0.360, yet it was placed fourth in the combined stability score. Again, the inclusion of any number of the genes provided a V score of <0.15, although the lowest V score was achieved with the inclusion of the three genes; *Lrpl14*, *Lube2* and *Lywhaz*. Of the three tissues, the embryo analyses yielded the best scores, followed by ovotestis. *Lef1a* was consistently found to be the least stable gene in all tissues. In comparison, NormFinder identified *Lhis2a* as the most stable gene in the embryo (stability value of 0.058; Table 2). However, the best combined pair was *Lacads* and *Lube2* (0.047). In the foot, *Lywhaz* was identified as the most stable gene (0.074) and was paired with *Lube2* (combined stability 0.066). In the ovotestis, *Lhis2a* was most stable (0.124), but the best combined pair of genes was *Lef1a* and *Lywhaz* (0.083), despite the fact that *Lef1a* presented the poorest (highest) individual gene stability value (0.243). As with geNorm, the embryo analyses yielded the best scores. *Lef1a* was found to be the least stable or second least stable individual gene in all tissues. In all analyses, the stability value of the best combined pair of genes was lower than that of any individual gene stability score.

**Table 2.**
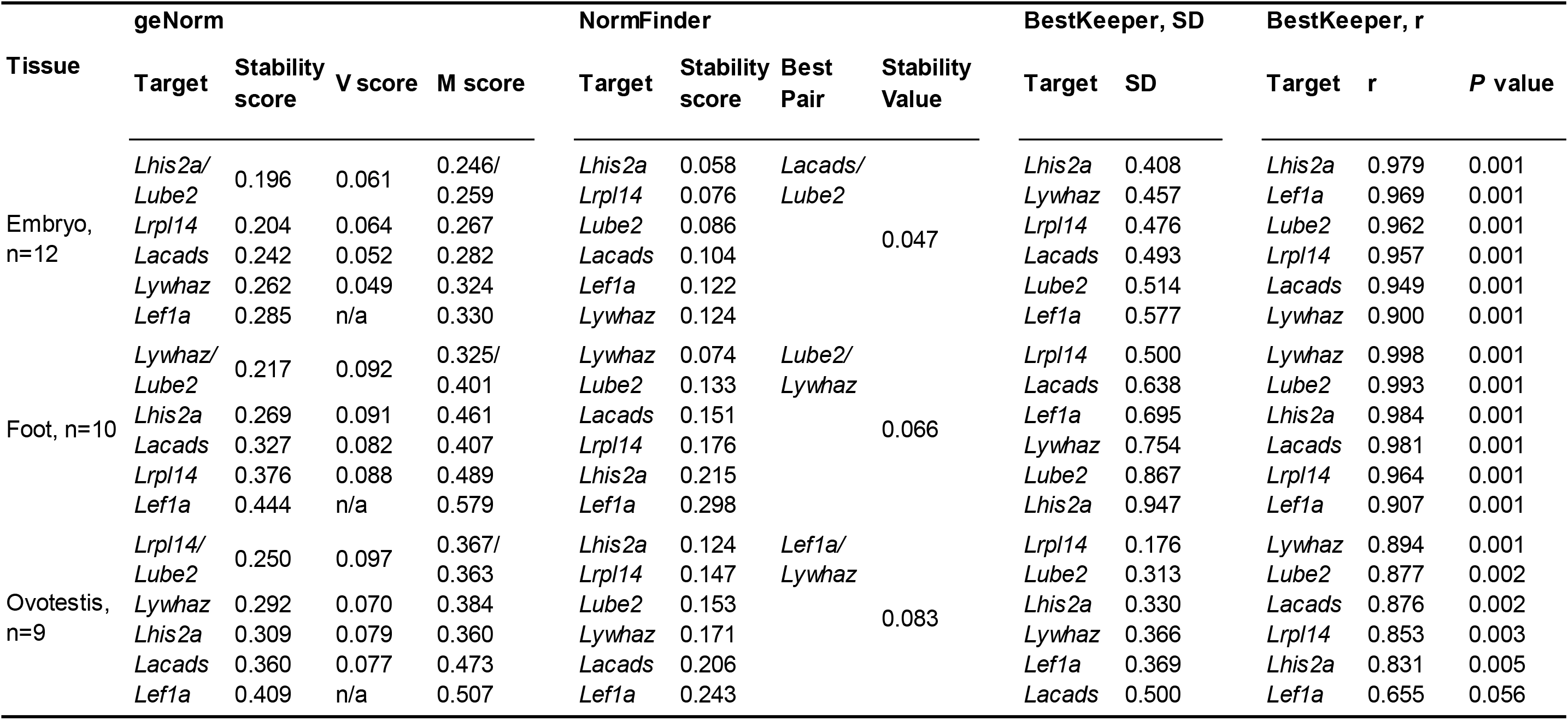
Gene expression stability results per tissue, using three different normalising control methods. geNorm provides the best paired combination of genes, with additional V scores indicating the best accumulative combination and individual M scores giving a measure of individual expression stability. Normfinder provides the best combined pair of genes with a separate associated stability score. Bestkeeper results are presented as both their correlation with the BestKeeper index (r), with associated probability values (*P*), and the standard deviation (SD) associated with the average Cq per gene.

The BestKeeper program provides two measures of gene stability, with a low SD and a high r value indicating a more stable control gene. In the embryo, the gene ranked as most stable according to SD was *Lhis2a* (0.408; Table 2), whereas the least stable gene was *Lef1a* (0.577). Every gene in the embryo analysis resulted in a highly significant positive correlation with the BestKeeper index (*P* = 0.001). *Lhis2a* demonstrated the highest correlation, with an r value of 0.979, and *Lywhaz* the lowest with an r value of 0.900. In the foot, *Lrpl14* was ranked as most stable according to SD (0.500), whereas the least stable gene was *Lhis2a* (0.947). With the exception of *Lef1a* in the ovotestis, every gene/tissue combination showed a significant correlation with the BK index.

### Relative expression of cytoskeletal genes

Relative expression of *Ldia2* transcripts depends upon the genetic background of the mother (Figure 1; Tables 3 and 4). Thus, levels of Ldia2 transcripts in embryos derived from a genetically sinistral mother *dd* were 0.006 (*Ldia2 3’UTR*, 0.6%) or 0.03 (*Ldia2 ORF*, 3%) relative to embryos from a wild-type *DD* mother; levels of the same transcripts in offspring of a heterozygote mother *Dd* were about half that of wild-type, 0.56 (*Ldia2 3’UTR*, 0.56%) or 0.48 (*Ldia2 ORF*, 0.48%). In comparison, there were few significant differences in the expression of the other genes, including *Ldia1*, with the exception of *Larp2/3-3* (*DD*:*Dd*, and *Dd*:*dd*) and *Larp2/3-1a* (*Dd*:*dd*). Notably, the relative differences in expression of *Ldia2* transcripts were much less striking in ovotestis, though still significantly lower, 0.81 (*Dd*) and 0.69 (*dd*) using *Ldia2 3’UTR* and 0.80 (*Dd*) and 0.62 (*dd*) using *Ldia2 ORF*; in foot tissue, there were no significant differences in expression between *Ldia2* transcripts from snails of different genotype (Figure 1; Tables 3 and 4).

**Table 3.** Normalised relative quantities (NRQ) of each gene, presented as a geometric mean per genotypic group (Geno), relative to different genotypes. Heterozygote snails, *Dd*, were not used with foot tissue.

|  | Genotype | N | *Larp2/3-1a* | *Larp2/3-3* | *Ldia1 3'UTR* | *Ldia2 3'UTR* | *Ldia2 ORF* | *Lfat* | *Lfry* |
| --- | --- | --- | --- | --- | --- | --- | --- | --- | --- |
| Embryo | *DD* | 6 | 1 | 1 | 1 | 1 | 1 | 1 | 1 |
|  | *Dd* | 5 | 1.059 | 0.735 | 0.969 | 0.563 | 0.476 | 1.052 | 0.984 |
|  | *dd* | 6 | 0.926 | 0.960 | 1.111 | 0.006 | 0.029 | 1.103 | 0.960 |
| Foot | *DD* | 5 | 1 | 1 | 1 | 1 | 1 | 1 | 1 |
|  | *dd* | 5 | 1.218 | 1.053 | 0.973 | 1.154 | 1.425 | 1.217 | 1.041 |
| Ovotestis | *DD* | 14 | 1 | 1 | 1 | 1 | 1 | 1 | 1 |
|  | *Dd* | 8 | 1.087 | 0.758 | 0.843 | 0.800 | 0.809 | 0.832 | 0.910 |
|  | *dd* | 14 | 0.965 | 0.877 | 1.017 | 0.619 | 0.686 | 0.847 | 0.878 |

**Table 4.**
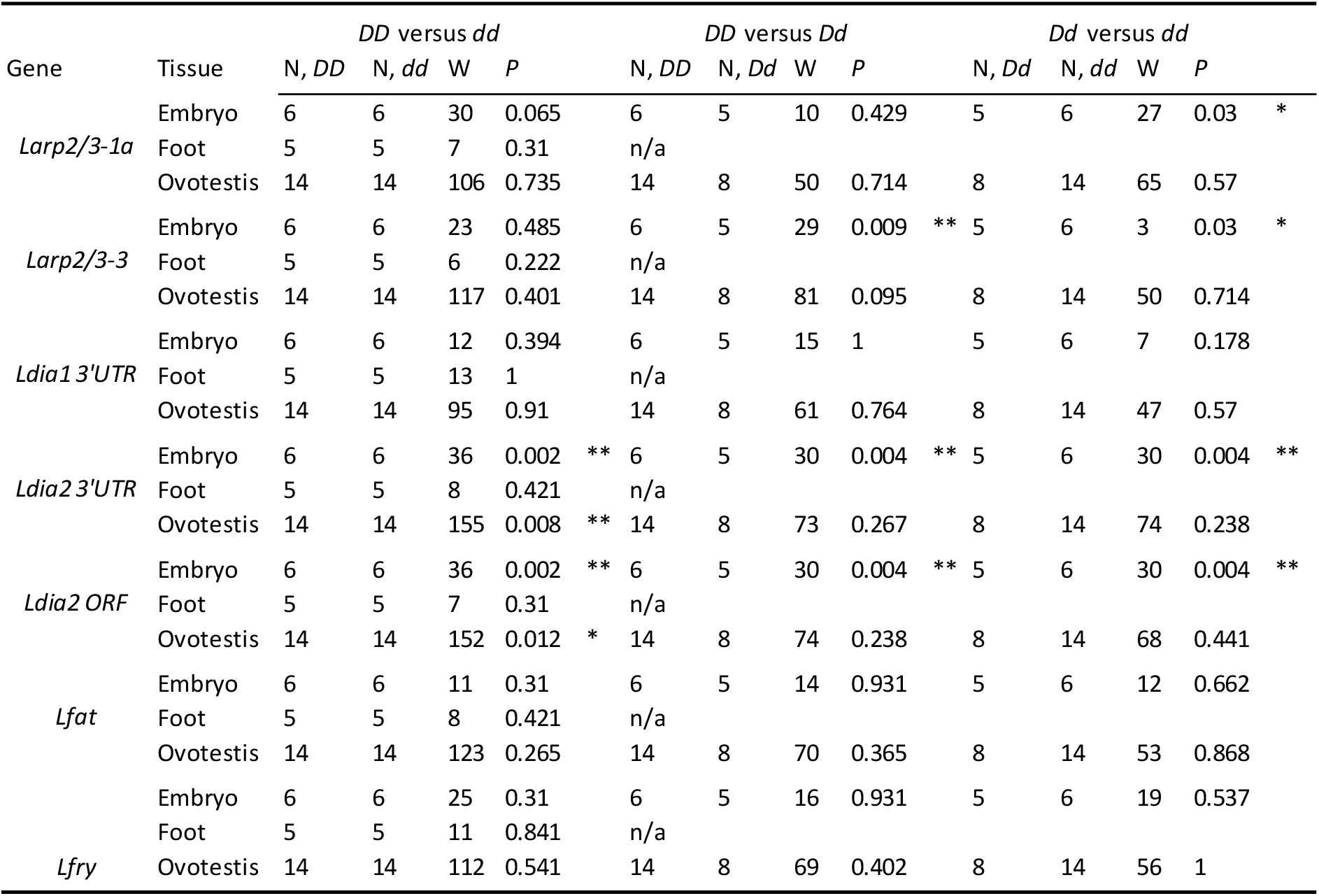
Wilcoxon rank test results for pairwise comparisons between genotypes *DD*, *Dd* and *dd* within embryo, foot and ovotestis tissue for cytoskeletal genes. The Wilcoxon rank value (W) is presented with the associated probability value (*P*). Statistical significance is highlighted via * <0.05, ** <0.01.

**Figure 1.**
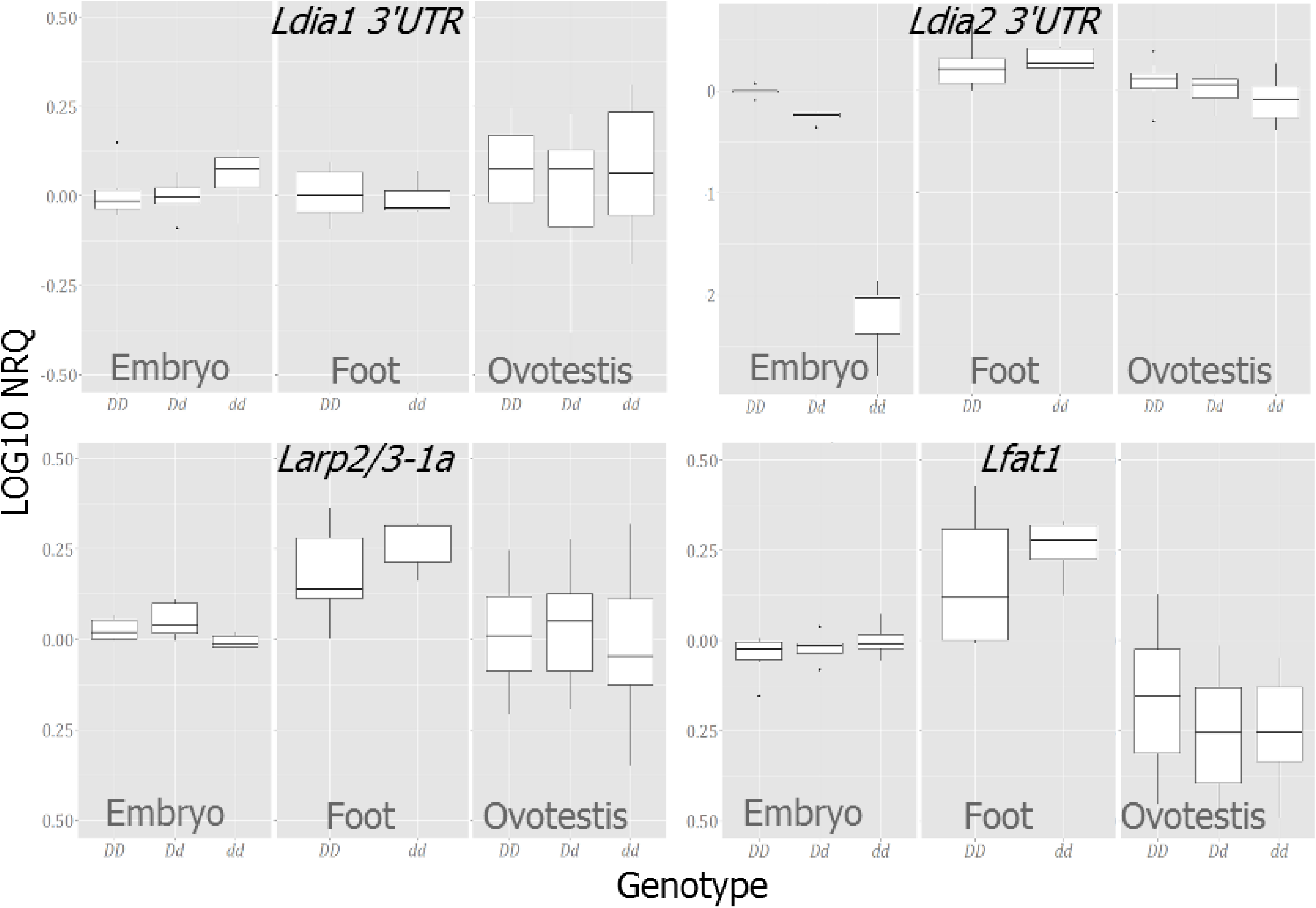
Boxplots showing Log scale NRQ values (LOG10 NRQ) for four different genes in in three genotypes, *DD*, *Dd* and *dd* across three tissues, embryo, foot and ovotestis tissue, relative to single standard. *Ldia2 3’UTR* transcripts are almost absent from the embryo in *dd* individuals, and also show reduced expression in the ovotestis. This effect is not seen in the foot tissue. The graphs also show that, in general, between-sample variation is least in the embryo.

All of the tested genes were relatively depleted in the single-cell embryo, relative to ovotestis and foot, except *Ldia2*, which was enriched (Figure 2; Table 5). *Larp*, *Lfat* and *Lfry* transcripts were reduced to ~0.03 to 0.27 of the level in embryo compared to ovotestis, and ~0.11 to 0.38 when comparing foot to single-cell embryo. In comparison, levels of *Ldia2* expression were ~1.27 to 2 times higher in the single-cell embryo compared to the ovotestis and ~2.8 times higher when compared to the foot tissue.

**Table 5.** Normalised relative quantities (NRQ) of each gene, presented as a geometric mean per genotypic group (Geno), relative to different tissues. Heterozygote snails, *Dd*, were not used with foot tissue.

| Tissue | Genotype | N | *Larp2/3-1a* | *Larp2/3-3* | *Ldia1 3'UTR* | *Ldia2 3'UTR* | *Ldia2 ORF* | *Lfat* | *Lfry* |
| --- | --- | --- | --- | --- | --- | --- | --- | --- | --- |
| Embryo | *DD* | 6 | 0.069 | 0.224 | 0.103 | 2.835 | 1.973 | 0.14 | 0.336 |
|  | *Dd* | 5 | 0.077 | 0.175 | 0.105 | 1.676 | 0.988 | 0.154 | 0.348 |
|  | *dd* | 6 | 0.065 | 0.219 | 0.118 | 0.019 | 0.058 | 0.158 | 0.33 |
| Foot | *DD* | 5 | 0.652 | 0.656 | 0.91 | 1.012 | 0.708 | 1.062 | 0.88 |
|  | *dd* | 5 | 0.784 | 0.682 | 0.875 | 1.153 | 0.997 | 1.277 | 0.905 |
| Ovotestis | *DD* | 14 | 1.935 | 1.068 | 1.049 | 1.418 | 1.553 | 1.339 | 1.239 |
|  | *Dd* | 8 | 2.256 | 0.869 | 0.95 | 1.218 | 1.348 | 1.196 | 1.21 |
|  | *dd* | 14 | 1.82 | 0.914 | 1.04 | 0.855 | 1.039 | 1.105 | 1.06 |

**Figure 2.**
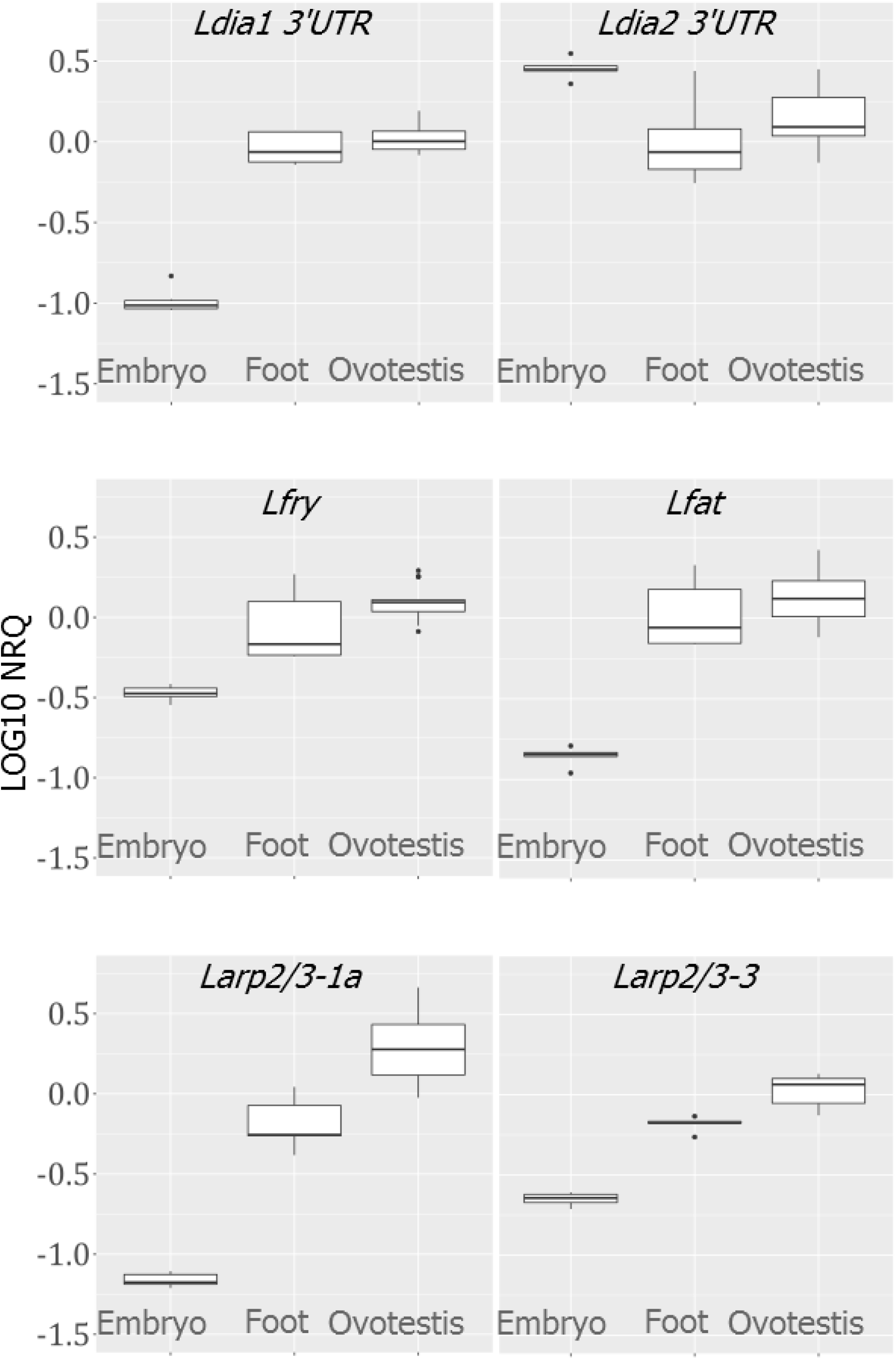
Boxplots showing Log scale NRQ values (LOG10 NRQ) for six different genes in three different tissue), using a single genotype *DD*, and relative to a single standard. The expression of five genes is depleted in the embryo, with the exception of *Ldia2*.

## DISCUSSION

Individually, all six gene targets were found to provide stable endogenous controls across all tissues, with the possible exception of *Lef1a*. However, the best individual gene and combination of genes differed between tissues used and analysis program. As it is recommended to use more than one control gene in combination in an experiment, then a tissue specific analysis is advisable prior to the experiment proper. Whether adding a third gene is worth the additional time and resources will depend on the individual experiment and the extent of the increase in stability gained.

For our analysis of gene expression in the early embryo, we used a combination of *Lhis2a*, *Lube2* and *Lyhwaz*/*Lrpl14*. A key finding was that transcripts of all genes except dextral-derived *Ldia2* were relatively depleted in the embryos (Figure 2). In comparison, the frameshifted version of *Ldia2* was severely depleted in the embryos (Figure 1), but these differences were less evident in the ovotestis and not evident in the foot tissue (Figure 2). The conclusion is that ability to detect the dynamics of nonsense mediated decay of RNA must therefore be highly dependent upon the tissue used.

### Genes to use as endogenous controls in different tissues

Within the embryo, all three algorithms ranked *Lhis2a* as the single most stable single gene, but there was less consensus for the rankings of the remaining endogenous controls. Generally, *Lhis2a*, *Lrpl14* and *Lube2* were in the top three most stable genes across software and tissue (Table 2). For the foot tissue analyses, due to the agreement of the different algorithms, *Lywhaz* and *Lube2* should be used as endogenous normalising controls. In comparison, the ovotestis results were more varied. The results of the geNorm analysis show that the use of *Lrpl14* and *Lube2* would provide an acceptable endogenous control measure, and the inclusion of a third gene, *Lywhaz*, indicates the most stable combination of genes.

*Lef1a* was consistently ranked least stable in all analyses of foot and ovotestis tissue, and often in the embryo, interesting because it has been a common choice by others as an endogenous control (Foster, Lukowiak & Henry, 2015; *van Nierop, et al., 2006*). However, we found that it is still acceptable for use, just not necessarily the gene of choice. The reason for the relatively poor performance may be due to a low level of expression rather than variable expression, indicated in the amplification efficiency experiments (Table 1). *Lef1a* may thus provide a reliable endogenous control gene when using an increased cDNA concentration.

Compared to the other tissues assessed, the embryo was found to be least variable (Figure 2). There are many reasons why some tissues may be more variable than others. In our experiments, it was difficult to temporally control the extraction of the ovotestis (e.g. time since egg-laying), and especially to make sure that it was free of contaminating hepatopancreas. In comparison, the embryos were from a clean and temporally controlled sample.

All three analytical programs used here provided a unique aspect of the data analysis. geNorm provided a measure of the optimum number of genes to include in the analysis and an advised cut-off value (V, <0.15) for an acceptable endogenous control gene combination. BestKeeper output a quotable measure of SD for each gene and a statistical measure of the relatedness of gene expression. Finally, NormFinder provided valuable information on the experimental design; calculating variation created both within and between experimental groups and importantly provides an alternative to pairwise comparison methods.

### Comparing expression in different tissues

Previously, we used qRT-PCR to show the formin *Ldia2* shows significant fold-change differences in the quantity of mRNA transcripts between different chirality-associated genotypes (Davison*, et al.*, 2016). Here we showed that the cytoskeletal genes, including *Ldia1*, are substantially depleted in the embryo, except for *Ldia2* (Figure 2). However, while the frameshifted version of *Ldia2* was severely depleted in single-cell embryos (Figure 1), these differences were less evident in the ovotestis and not evident in the foot tissue (Figure 2), indicating that the ability to detect nonsense-mediated decay must be tissue dependent.

As the cellular processes associated with variations in *L. stagnalis* chirality are predominantly cytoskeletal (Davison*, et al.*, 2016; Shibazaki*, et al.*, 2004; Tee *et al.*, 2015), this work further emphasises the potential pitfalls of using the commonly employed endogenous control genes, actin or tubulin, without adequate testing of their expression stability. Specifically, it also suggests that *Ldia1*/*Ldia2* have different roles during development, despite the close sequence similarity, and that *Ldia2* may be particularly critical in early development, given the relatively high levels of transcript present.

Comprehensive studies of nonsense-mediated decay have not been performed in molluscs. However in nonsense-mediated decay studies, from yeast to mammals, decay has been observed in both a 5’ to 3’ and 3’ to 5’ direction of the mRNA, originating from either the 3’ end or exon-exon boundaries (Karousis, Nasif & Mühlemann, 2016; Lykke-Andersen & Jensen, 2015). The variation in starting position of nonsense-mediated decay limits interpretation of the differences between the reduction *Ldia2* in the 3’ UTR and ORF. However, the frameshift in the sinistral *Ldia2* should be present in all tissues, and therefore the resulting nonsense-mediated decay would be expected to be evident in all tissues. The lack of significant quantitative differences in the foot tissue suggests that nonsense-mediated decay may be obscured in actively transcribing tissues. As transcription does not begin in *L. stagnalis* before the 8-cell stage (Liu*, et al.*, 2014), so a single-cell embryo only contains maternal mRNAs that are transcribed prior to oviposition; ovotestis is presumably enriched for this same material. Further experiments with later stage embryos would presumably confirm this hypothesis.

### Conclusions

It was established that any of the six genes would provide acceptable endogenous controls to standardise gene expression between chiral genotypes within any of the three different tissues, perhaps with the exception of *Lef1a*. These primers should therefore permit rapid verification of endogenous controls suitable for use in qRT-PCR experiments assessing ovotestis, foot and embryo tissue within and between chiral variants of *L. stagnalis*, which was lacking previously.

## SUPPORTING INFORMATION

**Supplementary Information 1.** Fasta format alignments of genomic sequences against transcriptome and primer sequences for endogenous control genes and cytoskeletal genes.

**Supplementary Information 2.** Raw data for the qRT-PCR experiments.

## ACKNOWLEDGEMENTS

The authors would like to thank Maureen Liu and Sheila Keeble for assistance with some research planning and laboratory work. This work was mainly funded by a BBSRC DTP studentship to Harriet Johnson.

